# RGI-GOLVEN signalling promotes FLS2 abundance to regulate plant immunity

**DOI:** 10.1101/2021.01.29.428839

**Authors:** Martin Stegmann, Patricia Zecua-Ramirez, Christina Ludwig, Ho-Seok Lee, Brenda Peterson, Zachary L. Nimchuk, Youssef Belkhadir, Ralph Hückelhoven

## Abstract

Endogenous plant signalling peptides regulate developmental and growth-related processes. Recent research indicates that some of these peptides are classified as phytocytokines as they have regulatory functions during plant immune responses. However, the mechanistic basis for phytocytokine-mediated immune modulation remains largely elusive. Here, we identify GOLVEN2 (GLV2) peptides as novel phytocytokines in *Arabidopsis thaliana*. By peptide application, precursor overexpression and loss-of-function studies we show that GLV2 enhances sensitivity of plants to elicitation with the bacterial flagellin epitope flg22. GLV2 is perceived by ROOT MERISTEM GROWTH FACTOR 1 INSENSITIVE (RGI) receptors and RGI3 forms an flg22-induced complex with the flg22-receptor FLAGELLIN SENSITIVE 2, suggesting that RGIs are part of activated pattern recognition receptor signalling platforms. GLV2 perception increases posttranscriptional FLS2 abundance and RGIs promote FLS2 protein accumulation. Thus, GLV-RGI signalling controls above ground plant immunity via a novel mechanism of phytocytokine activity.

## Introduction

Receptor kinases (RKs) sense external and internal cues, including plant peptide hormones, to regulate a diverse range of plant physiological responses, ranging from growth/development to plant immunity and stress responses. RKs can serve as pattern recognition receptors (PRRs) for detecting microbe-associated molecular patterns (MAMPs) to trigger pattern-triggered immunity (PTI), the first line of induced defence of plants against invading microbes. A well characterized example is the perception of bacterial flagellin by the leucine-rich repeat (LRR)-RK FLAGELLIN-SENSITIVE 2 (FLS2) (Gómez-Gómez and Boller, 2000; Zipfel et al., 2004). FLS2 binds to a 22 amino acid peptide derived from bacterial flagellin (flg22) which is released by host hydrolytic enzymes (Buscaill et al., 2019). Perception of flg22 results in the activation of downstream signalling culminating in basal resistance against pathogens (Couto and Zipfel, 2016).

In addition to MAMPs, plants can also perceive endogenous peptides to sense danger and modulate immune signalling by PRRs. Research in recent years has identified several classes of such immunomodulatory peptides. These are functionally analogous to metazoan cytokines and thus referred to as phytocytokines (Gust et al., 2017). Importantly, mammalian cytokine and MAMP signalling share downstream signalling pathways and regulate each other. However, the modulatory actions of phytocytokines on plant immune responses remain for the most part poorly characterized (Gust et al., 2017; Trinchieri and Sher, 2007). Among the most prominent plant immune amplifying peptides are the PLANT ELICITOR PEPTIDES (PEPs) which derive from the C-terminus of PROPEP precursors and are proteolytically released by METACASPASE 4 (Bartels et al., 2013; Hander et al., 2019). *PROPEP* gene expression is triggered by biotic stresses, including microbial infection and MAMP perception (Bartels and Boller, 2015). Two additional peptides that work in a similar fashion are PATHOGEN-ASSOCIATED MOLECULAR PATTERN (PAMP)-INDUCED PEPTIDES (PIPs) and the recently discovered SERINE-RICH ENDOGENOUS PEPTIDES (SCOOPs) (Gully et al., 2019; Hou et al., 2014). PIPs and SCOOPs are also implicated in feed-forward loops to boost immune stimulation during infections.

An important family of plant peptides regulating growth and development are RAPID ALKALINISATION FACTOR (RALF) peptides, which control growth by inhibiting cell elongation through apoplastic pH modulation (Morato do Canto et al., 2014). RALFs are also involved in plant fertilization where they control pollen tube integrity (Ge et al., 2017; Mecchia et al., 2017). In *Arabidopsis thaliana* (hereafter referred to as Arabidopsis), some RALF peptides are perceived by a FERONIA (FER)-LORELEI-LIKE GPI-ANCHORED PROTEIN 1 (LLG1) heterocomplex (Haruta et al., 2014; Stegmann et al., 2017; Xiao et al., 2019). FER-LLG1 controls PTI by serving as a RALF-regulated scaffold for the assembly of PRR complexes at the plasma membrane (Stegmann et al., 2017; Xiao et al., 2019). PHYTOSULFOKINE (PSK) and PLANT PEPTIDE CONTAINING SULFATED TYROSINE 1 (PSY1) are tyrosine-sulfated peptides that promote growth through cell elongation via partially overlapping pathways (Matsubayashi, 2014; Sauter, 2015). PSK is perceived by PSK RECEPTOR 1 (PSKR1) and PSKR2, while recognition of PSY1 involves PSY1-RECEPTOR (PSYR) (Matsubayashi, 2014). Overexpression of PSK precursors and mutating *PSKR1* or *PSY1R* enhances susceptibility to hemibiotrophic bacteria and resistance to necrotrophic fungi (Igarashi et al., 2012; Mosher et al., 2013), while exogenous PSK treatment inhibits late MAMP-triggered responses. However, the molecular mechanism of signalling cross-talk between these peptide pathways and PTI remains largely unknown (Segonzac and Monaghan, 2019).

GOLVEN (GLV) peptides, which are also known as ROOT MERISTEM GROWTH FACTOR (RGF) or CLAVATA 3/EMBRYO SURROUNDING REGION-LIKE (CLEL) peptides are encoded by a family of 11 genes in Arabidopsis (Fernandez et al., 2013). Hereafter, we will refer to these peptides as GLVs. They are produced as preproproteins and require posttranslational modification steps for maturation and extracellular release. Fully maturated GLV peptides are usually 13-18 amino acids in length. The GLV1 and GLV2 peptide precursors are processed by SUBTILASE (SBT)6.1/SITE-1 PROTEASE (S1P) and SBT3.8 on their journey through the secretory pathway (Ghorbani et al., 2016; Stührwohldt et al., 2020). For full activity, GLVs are sulfated on a conserved tyrosine residue targeted by TYROSYLPROTEIN SULFOTRANFERASE (TPST) (Matsuzaki et al., 2010; Whitford et al., 2012). GLVs play multiple roles on root development and they have been associated with root gravitropism, root stem cell niche maintenance and lateral root formation (Fernandez et al., 2015, 2020; Matsuzaki et al., 2010; Meng et al., 2012; Whitford et al., 2012). Several *GLV* genes can be simultaneously expressed in individual cell types or organs, suggesting redundant functions. For example, 7 of the 11 *GLV* members are expressed in the root apical meristem (Fernandez et al., 2013; Matsuzaki et al., 2010) and simultaneous knockout of the three *GLV* genes *GLV11*, *GLV5* and *GLV7* results in a short root phenotype. GLV1 and GLV2 regulate root gravitropism (Whitford et al., 2012) and both GLV6 and GLV10 repress asymmetric cell divisions during lateral root initiation (Fernandez et al., 2020).

GLV peptides are perceived by a family of five LRR-RKs that function during lateral root initiation and root stem cell niche maintenance. In the latter context, three groups independently found RGF1 INSENSITIVE (RGI) 1-5 to perceive GLV11 (Shinohara et al., 2016; Ou et al., 2016; Song et al., 2016). A *rgi1/2/3* mutant has a stunted primary root and is insensitive to GLV11 treatment (Shinohara et al., 2016). The short root phenotype of *rgi1/2/3* is further enhanced when *RGI4* and *RGI5* are genetically eliminated (Ou et al., 2016). GLV11 binds to the ectodomain of RGI1, RGI2 and RGI3 (Shinohara et al., 2016; Song et al., 2016). The crystal structure of the RGI1-GLV11 complex revealed that a conserved RxGG motif is required to bind to the sulfated tyrosine of GLV11 (Song et al., 2016). This motif is conserved among all 5 RGIs, suggesting that they share a similar recognition mechanism for related GLV peptides. Consistently, RGI1 was able to bind to GLV5, GLV7 and GLV10 *in vitro*, similar as to GLV11 (Song et al., 2016). Genetically, *RGI1* and *RGI2* are the major genes implicated in root meristematic activity whereas the role of *RGI3* in this process is less pronounced (Shinohara et al., 2016). Consistent with a predominant role for *RGI1* and *RGI2*, expression of *RGI2* under the control of its native promoter largely rescues the short root phenotype of the *rgi1/2/3/4/5* quintuple mutant (Ou et al., 2016). During GLV-mediated inhibition of lateral root initiation, RGI1, RGI4 and RGI5 are the main receptors for GLV6 and GLV10. GLV10-induced inhibition of lateral root emergence is abolished in a *rgi1/4/5* triple mutant (Fernandez et al., 2020).

RGI1 can form a GLV11-induced complex with members of the SOMATIC EMBRYOGENESIS RECEPTOR KINASE (SERK) family (Song et al., 2016) which are critical co-receptor proteins for a multitude of LRR-type RKs (Ma et al., 2016). Consistently, higher order *serk* mutants are insensitive to the root growth promoting effect of GLV11 (Song et al., 2016). To relay the extracellular signal into a cellular response, MITOGEN-ACTIVATED PROTEIN KINASES (MAPKs) become activated after GLV receptor complex activation. MAPK6 functions downstream of RGI1/3/4 in restricting lateral root emergence. A mutation in MAPK6 can restore the formation of lateral roots in a conditional GLV6 overexpression line (Fernandez et al., 2020). MPK6, together with MPK3 also acts downstream of GLV11 perception by RGI1-5 during the control of meristematic activity. Here, MPK3 and MPK6 are activated by the MAPKKK YODA and the MAPKK MKK4 and MKK5 (Lu et al., 2020; Shao et al., 2020). These studies indicate that shared pathways operate downstream of RGIs in different physiological contexts.

In the primary root, GLV11 controls meristematic activity through stimulating the accumulation of the transcription factor PLETHORA (PLT) both transcriptionally and posttranscriptionally (Matsuzaki et al., 2010). GLV11 regulates PLT stability through RGI-dependent transcriptional upregulation of the transcription factor RGF1-INDUCIBLE TRANSCRIPTION FACTOR 1 (RITF1) and subsequent distribution of reactive oxygen species (ROS) along the developmental zones of roots (Yamada et al., 2019). To control root gravitropism, GLV1 and GLV3 increase the abundance and subcellular trafficking of the auxin efflux carrier PINOID 2 (PIN2) largely independent of transcription, indicating that GLV signalling can influence the accumulation and/or stability of diverse proteins and, in the case of plasma membrane proteins, can additionally regulate their endocytosis, vesicle delivery and/or endosomal sorting (Whitford et al., 2012). Yet, a mechanistic link from RGI receptors to PIN2 remains elusive.

A multitude of specific functions of GLV peptides are known for several aspects of root development. A most recent report also indicates a function for the leaf expressed GLV4 in plant immunity. *GLV4* gene expression is rapidly induced in response to flg22 treatment and inducible expression of GLV4 peptide precursors induces PTI-like responses dependent on leaf-expressed *RGI4* and *RGI5* (Wang et al., 2021). These findings suggest an involvement of GLV peptides in the regulation of above ground immunity. Yet, it remains unclear whether GLV peptides modulate immune signalling by PRRs and if so, what would be the underlying molecular mode-of action.

Here, we show that additional shoot expressed GLV peptides have a novel function in modulating immune responses. Overexpression of *GLV1* and *GLV2* peptide precursor genes, as well as simultaneous genetic loss of leaf expressed *GLVs* shows that GLV peptides are positive regulators of flg22-triggered PTI. GLV2 is perceived by leaf expressed RGI3 which forms a flg22-induced complex with FLS2, suggesting that GLV-RGI signalling constitutes a novel peptide-receptor pathway controlling flg22-triggered PTI. GLV2 perception increases FLS2 abundance, providing a mechanistic basis for GLV-mediated PTI regulation and a novel role for GLV peptides in above ground tissue as FLS2 signalling-promoting phytocytokines.

## Results

### *GLVs* are involved in flg22-triggered PTI

Several endogenous peptide-coding genes associated with immunity are transcriptionally regulated in response to pathogen infections or elicitor perception (Gully et al., 2019; Igarashi et al., 2012; Mosher et al., 2013). To identify novel endogenous peptides involved in immunity we mined publicly available RNAseq datasets. We focused on members of peptide families that displayed differential transcriptional regulation during biotic stress. With this approach, we found that *GLV1* and *GLV2*, two genes that code for peptides involved in root gravitropism, are transcriptionally downregulated in leaves after infection with the virulent bacterial strains *Pseudomonas syringae* pv. *maculicola* (*Psm*) and *Pseudomonas syringae* pv. *tomato* (*Pto)* DC3000, albeit with different strengths (Figure 1 – figure supplement 1)(Bernsdorff et al., 2016; Howard et al., 2013). Notably, *GLV1* and *GLV2* did not show altered expression when flg22 was used alone. Based on these observations, we hypothesized that *GLV1/2* can influence bacterial colonization and perhaps flg22-triggered responses. To address this, we first used flg22-induced ROS and seedling growth inhibition (SGI) assays to measure FLS2 activation in *GLV1* and *GLV2* overexpression lines (*35S::GLV1* and *35S::GLV2*) (Whitford et al., 2012). Both response outputs were enhanced in *35S::GLV1* and *35S::GLV2* lines (Figure 1A, B). Also, *GLV1* and *GLV2* overexpression lines displayed enhanced resistance against the moderately virulent bacterial pathogen *Pto* DC3000 lacking the effector molecule coronatin (*Pto* DC3000 COR-) (Figure 1C). This demonstrates that overexpression of *GLV1* and *GLV2* elevates flg22-triggered PTI against bacterial pathogens.

**Figure 1:**
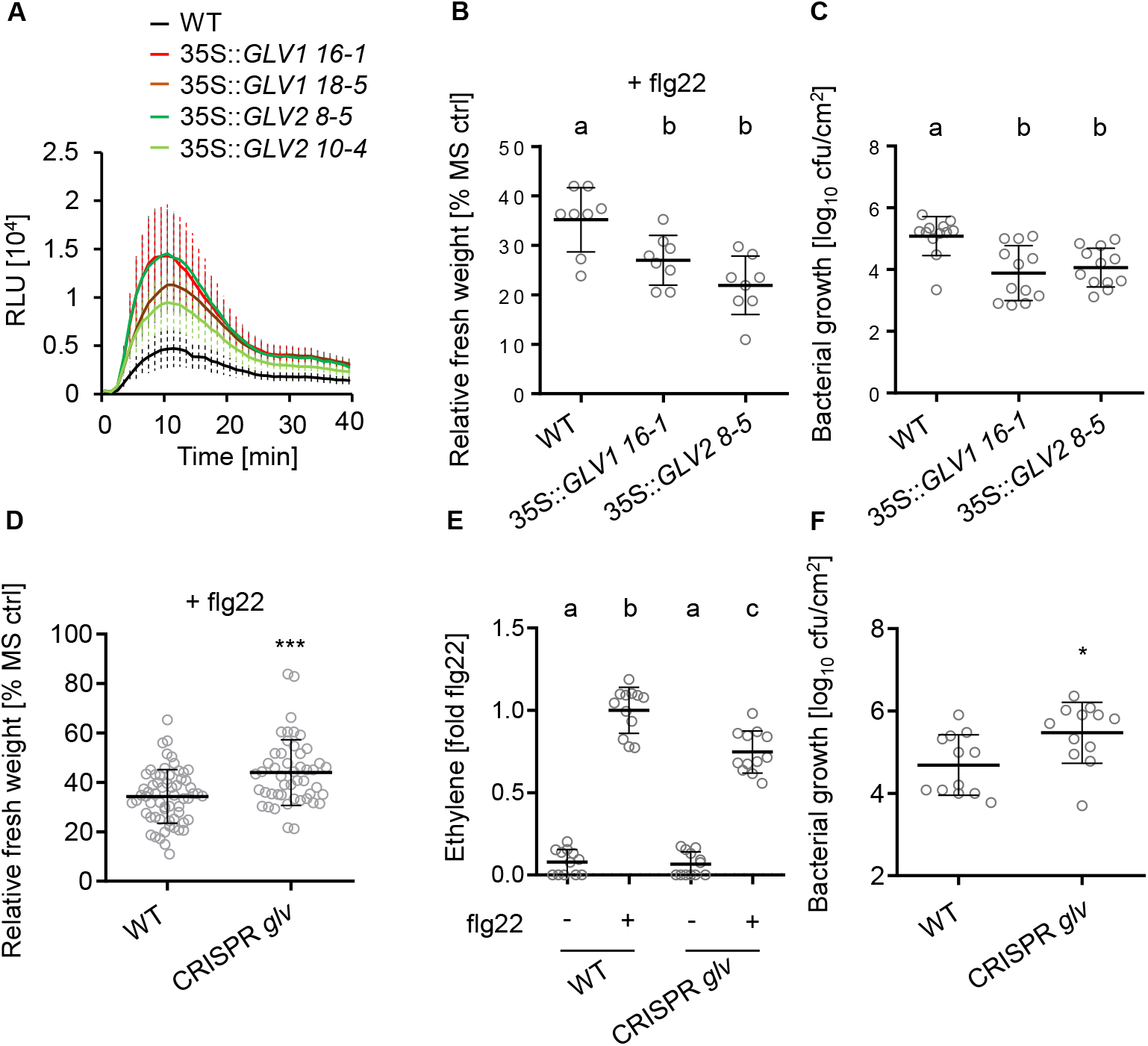
*GLVs* are positive regulators of flg22-triggered PTI. (A) ROS burst in wild type (WT) and *GLV1* and *GLV2* overexpression lines after elicitation with 100 nM flg22. The kinetic of ROS production is shown over 40 min −/+ SE, n=8. RLU = Relative light units. *GLV1* and *GLV2* overexpression lines show increased flg22-induced ROS production (B) 5 day-old seedlings of WT, *35S::GLV1 16-1* and *35S::GLV2 8-5* were treated with 100 nM flg22 for 7 days before measuring fresh weight. Shown is the mean of relative fresh weight compared to the MS medium control −/+ SD, n=8 (one-way ANOVA, Dunnett post-hoc test, a-b p<0.05). The *GLV1* and *GLV2* overexpression lines are more sensitive to flg22-induced SGI (C) Mean colony-forming units (cfu) of *Pto* DC3000 COR-4 days after spray infection of WT, *35S::GLV1 16-1* and *35S::GLV2 8-5*; n=12 from 3 pooled biological replicates −/+ SD (one-way ANOVA, Dunnett post-hoc test; a-b, p<0.01). The *GLV1* and *GLV2* overexpression lines are more resistant to infection (D) 5-day-old seedlings of Col-0^Cas9^ WT and CRISPR *glv* were treated with 100 nM flg22 for 7 days before measuring fresh weight. Shown is the mean of relative fresh weight compared to the MS medium control −/+ SD, n=40 from 5 pooled biological replicates (Students T-test, ***p<0.001). The CRISPR *glv* line is less sensitive to flg22-induced SGI (E) Ethylene production was measured in Col-0^Cas9^ WT and CRISPR *glv* 3.5 h after elicitation with 500 nM flg22. Shown is the mean of n=12 from 3 independent biological replicates −/+ SD (one-way ANOVA, Tukey post-hoc test, a-b, p<0.001; a-c, p<0.001; b-c, p<0.001). Ethylene production was normalized to the average of WT responses to flg22 set as 1. The CRISPR *glv* line shows reduced ethylene accumulation compared to the WT (F) Mean cfu of *Pto* DC3000 COR-4 days after spray inoculation of Col-0^Cas9^ WT and CRISPR *glv*; n=12 from three pooled biological replicates −/+ SD (Students T-Test, *p<0.05). The CRISPR *glv* line is more susceptible compared to the WT. Similar results were obtained in at least 3 independent experiments.

Then, we confirmed by targeted proteomics on apoplastic wash fluids that *GLV2* overexpression results in an effective increase of mature GLV2 peptide in the extracellular space of leaves. We specifically searched for the tyrosine-sulfated and proline-hydroxylated GLV2 [DMD(TyrSO_3_H_2_)NSANKKR(Hyp)IHN] that was previously identified in root exudates of *GLV2* overexpression lines (Whitford et al., 2012). GLV2 peptide accumulation was strongly increased in *35S::GLV2* plants when compared to wild type (Figure 1 – figure supplement 2). Next, we analysed whether the abundance of mature GLV2 changes upon elicitation with flg22 and only observed a mild but not significant reduction in both the wild type and the overexpression line. Thus, we propose that GLV2 precursor gene expression and secretion is constitutive under the conditions tested and not influenced by flg22-dependent immune activation.

Next, we studied the function of GLV peptides in flg22-triggered PTI by analysing loss of function mutants. First, we tested whether flg22-triggered responses and resistance to *Pto* DC3000 COR-is affected in a *glv2-1 amirGLV1* double mutant line (Whitford et al., 2012). We noticed a mild reduction in sensitivity towards flg22 in SGI experiments (Figure 1 – figure supplement 3A), but only observed a trend towards increased susceptibility upon *Pto* DC3000 COR-infections (Figurer 1 – figure supplement 3B). This suggests that additional GLV peptides may be important for flg22-triggered PTI in leaf tissue, similar to GLV-regulated developmental responses in roots (Fernandez et al., 2020; Matsuzaki et al., 2010; Whitford et al., 2012). Consistent with this, the recently identified GLV4 immune-inducing peptide and 4 additional GLV members are expressed in shoot and leaf tissue (Fernandez et al., 2013; Wang et al., 2021). Thus, we tested a higher order CRISPR-Cas9 *glv* mutant in which the majority of leaf expressed *GLVs* (*GLV1*, *GLV2*, *GLV6*, *GLV7*, *GLV8*, *GLV10*) and the related *CLE18* gene are mutated (Fernandez et al., 2020). The resulting mutant (CRISPR *glv*), showed reduced flg22-triggered ethylene production and SGI, respectively (Figure 1D, E). The CRISPR *glv* mutant was also significantly more susceptible to *Pto* DC3000 COR-(Figure 1F). Together, the data indicates that GLVs are key determinants modulating the intensity of flg22-triggered PTI.

### GLV2 peptides increase sensitivity of plants to flg22 perception

Next, we obtained synthetic GLV2 peptides by chemical synthesis and tested whether the GLV2 precursor-derived peptide can also influence flg22-triggered immunity. GLV2 itself did not induce classical PTI outputs such as an ROS burst response (Figure 2A). However, treating leaf discs with GLV2 together with flg22 increased flg22-triggered ROS production, a response induced a few minutes after elicitor application (Figure 2B). This effect was even stronger when leaf discs were pre-treated for five hours prior to flg22 addition (Figure 2B), suggesting that GLV2 perception results in increased sensitivity of plants to subsequent immune stimulation by flg22. Consistent with that, longer term co-treatment promoted flg22-triggered signalling outputs. Co-treatment between flg22 and GLV2 increased flg22-triggered ethylene production, *PR1* expression and SGI (Figure 2C, D, E). Finally, GLV2 treatment induced resistance to subsequent *Pto* DC3000 infection (Figure 2F). These findings reinforce the notion that exogenous application of GLV2 shows a similar effect on flg22-triggered PTI as endogenous overexpression of the peptide precursor gene.

**Figure 2:**
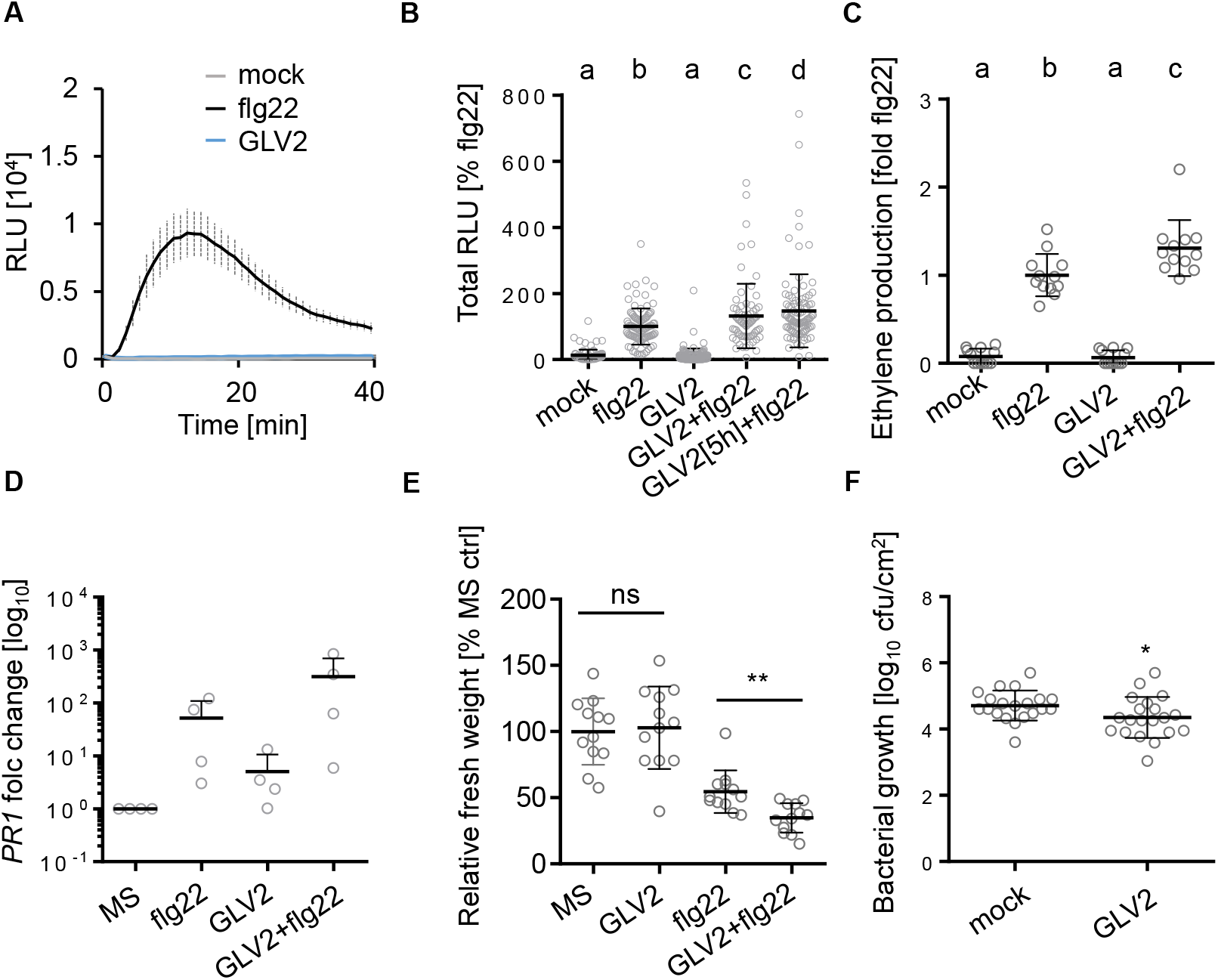
GLV2 treatment enhances flg22-triggered responses. (A) ROS burst in WT after elicitation with 100 nM flg22 or 1 μM GLV2. Shown is the kinetic of ROS production over 40 min −/+ SEM, n=8. RLU = Relative light units. GLV2 does not induce a classical ROS burst response. (B) ROS burst in WT after elicitation with 10 nM flg22, 1 μM GLV2, co-treatment or pre-treatment with 1 μM GLV2 for 5 h prior to elicitation with flg22. Shown is the mean of total ROS production (Total RLU) over 40 minutes normalized to the flg22 response set as 100% −/+ SD, n=96-120 of 15 pooled biological repeats (one-way ANOVA, Tukey post-hoc test, a-b/c/d, p<0.001; b-c, p<0.05; b-d, p<0.001). GLV2 co-treatment enhances flg22-triggered ROS production, which is further increased after pre-treatment for 5 h (C) Ethylene production was measured in WT after elicitation with 500 nM flg22, 1 μM GLV2 or co-treatment for 3.5 h. Ethylene production was normalized to the average flg22 response set as 1. Shown is the mean of n=12 from 3 independent biological replicates −/+ SD (one-way ANOVA, Tukey post-hoc test, a-b/c p<0.001; b-c, p<0.01). Ethylene production was normalized to flg22 treatment set as 1. GLV2 co-treatment increases flg22-triggered ethylene production (D) *PR1* expression was measured in 10-day-old WT seedlings treated for 24 h with 100 nM flg22, 1 μM GLV2 or co-treatment. Expression was normalized to the *UBQ* house-keeping gene. Shown is the mean −/+ SD of 4 independent biological repeats. GLV2 co-treatment increases flg22-induced *PR1* expression (E) 5-day-old WT seedlings were treated with 10 nM flg22, 1 μM GLV2 or co-treated for 7 days before measuring fresh weight. Shown is the mean of relative fresh weight compared to the MS medium control −/+ SD, n=12 (Students T-test, ns = not significant, **p<0.01). GLV2 does not induce SGI but co-treatment increases flg22-induced SGI. (F) Mean cfu of *Pto* DC3000 2 days after syringe inoculation and pre-treatment with 1 μM GLV2; n=20 from 5 pooled biological replicates −/+ SD (Students T-test, *p <0.05). GLV2 treatment induces resistance to *Pto* DC3000. Similar results were obtained in at least 3 independent experiments.

GLV2 requires sulfation at a conserved tyrosine residue by TPST to control meristematic activity and root gravitropism (Matsuzaki et al., 2010; Stührwohldt et al., 2020). The *tpst* mutants show increased immune responses which was mainly linked to PSK and PSY1-mediated immune regulation, both of which are also substrates of TPST (Igarashi et al., 2012; Mosher et al., 2013). Based on these previous findings we hypothesized that TPST-catalysed GLV tyrosine-sulfation may not be required for PTI. Surprisingly though, a GLV2 peptide mutant without sulfated tyrosine (GLV2^−S^) was ineffective and did not support flg22-triggered SGI (Figure 2 – figure supplement 1), suggesting that tyrosine sulfation is also critical for GLV2`s function during FLS2-triggered immunity.

### RGI receptor kinases are required for FLS2-mediated signalling

RGI1-5 are known receptors for GLV peptides (Shinohara et al., 2016; Ou et al., 2016; Song et al., 2016, Fernandez et al., 2020). Thus, we tested whether RGI1-5 may be involved in flg22-triggered responses. The *rgi1/2/3/4/5* quintuple mutant (hereafter referred to as *rgi5x*) showed reduced flg22-triggered ROS production (Figure 3A) and ethylene accumulation, respectively (Figure 3B). Because both assays are performed with leaf discs, these findings suggest that RGIs have additional functions in leaves where they are involved in flg22-triggered signalling. The *rgi5x* mutant also showed reduced sensitivity to flg22 in SGI experiments (Figure 3C), confirming a positive regulatory role of RGIs for flg22-induced responses. Consistent with the recently proposed role for leaf expressed RGIs in antibacterial resistance, the *rgi5x* mutant was more susceptible to *Pto* DC3000 COR-(Figure 3D) (Wang et al., 2021). The *rgi5x* mutant shows severely reduced root growth but no morphological phenotypes in above ground tissue (Ou et al., 2016). *RGI3* is one of three *RGIs* (*RGI3*, *RGI4, RGI5*) expressed in leaf tissue (Klepikova et al., 2016). We confirmed that *RGI3* is expressed in leaves from 5–6-week-old plants by semi-quantitative reverse transcription PCR (Figure 3 – figure supplement 1A). *RGI3* transcript accumulation was similar to *RGI4* but stronger than *RGI5*. This is in contrast to previous findings which analysed *RGI3* expression in shoots from seedlings (Wang et al., 2021). Thus, we attempted to complement the PTI phenotypes of *rgi5x* with *RGI3* driven under its native promoter (*pRGI3::RGI3*). Expression of *RGI3* complemented the reduced flg22-induced ethylene production phenotype of the *rgi5x* mutant (Figure 3E, Figure 3 – figure supplement 1B). Interestingly, the short root phenotype of *rgi5x* was not complemented by *pRGI3::RGI3* (Figure 3 – figure supplement 2A, B). Based on this, we propose that RGIs are selectively required in disparate processes such as root growth regulation and immunomodulation in leaves.

**Figure 3:**
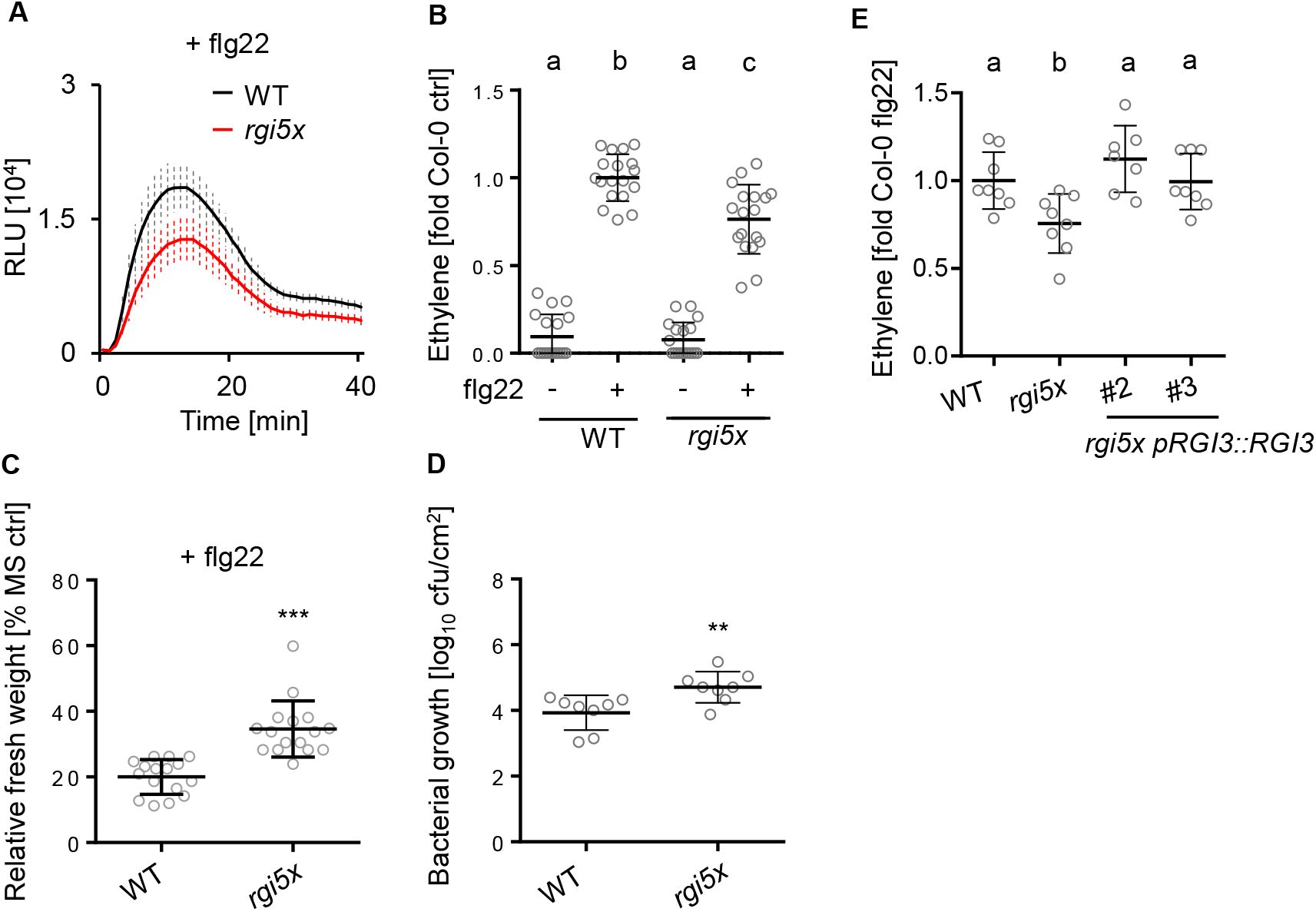
RGI receptors are involved in flg22-triggered PTI. (A) ROS burst in WT and *rgi5x* after elicitation with 100 nM flg22. Shown is the kinetic of ROS production over 40 min −/+ SE, n=8. RLU = Relative light units. The *rgi5x* mutant shows reduced flg22-triggered ROS production. (B) Ethylene production was measured in the indicated genotypes 3.5 h after elicitation with mock or 500 nM flg22. Ethylene accumulation was normalized to the average of WT responses to flg22 set as 1. Shown is the mean of n=18 from 3 independent biological replicates −/+ SD (one-way ANOVA, Tukey post-hoc test, a-b p<0.001; a-c, p<0.001; b-c, p<0.001). The *rgi5x* mutant shows reduced flg22-triggered ethylene production compared to the WT. (C) 5-day-old seedlings of WT and *rgi5x* were treated with 100 nM flg22 for 7 days before measuring fresh weight. Shown is the mean of relative fresh weight compared to the MS medium control −/+ SD, n=16 (Students T-test, ***p<0.001). The *rgi5x* mutant is less sensitive to flg22 (D) Mean colony-forming units (cfu) of *Pto* DC3000 COR-3 days after spray infection of the indicated genotypes; n=8 from two pooled biological replicates −/+ SD (Students T-test, **p<0.01). The *rgi5x* mutant is more susceptible to infection compared to the WT. (E) Ethylene production was measured in WT and *rgi5x* and *rgi5x pRGI3::RGI3 #2* and *#3* 3.5 h after elicitation with 500 nM flg22. Ethylene production was normalized to the average of WT responses to flg22 set as 1. Shown is the mean of n=8 from 2 independent biological replicates −/+ SD (one-way ANOVA, Dunnett post-hoc test, a-b p<0.05). Expression of native promoter-driven *RGI3* complements the flg22-induced ethylene phenotype of the *rgi5x* mutant. Similar results were obtained in at least 3 independent experiments.

### RGI3 perceives GLV2 to regulate FLS2-triggered PTI

Next, we tested whether RGIs are required for GLV2 function during PTI. We found that the GLV2-mediated increase in flg22-induced SGI and ethylene production was abolished in *rgi5x* mutants (Figure 4A, B). Therefore, RGIs are genetically required for the GLV2-triggered modulation of immune responses in leaf tissue. Since expression of *RGI3* alone is sufficient to rescue the GLV2-mediated increase in flg22-induced SGI in *rgi5x* mutants (Figure 4C), we predicted that RGI3 could act as a GLV2 receptor. Using *in vitro* pull downs, we demonstrated that a biotinylated GLV2 peptide (Biotin-GLV2), but not Biotin-flg22, physically interacted with the ectodomain of RGI3 (RGI3^ECD^) expressed from *Drosophila* cell culture (Figure 4D). Binding could be outcompeted with a 100-fold excess of unlabelled GLV2, but not with flg22. Conversely, Biotin-flg22, but not Biotin-GLV2, interacted with the FLS2^ECD^. Thus, RGI3 is a bona fide receptor for GLV2 and our data provide evidence for direct binding of GLV2 to an RGI.

**Figure 4:**
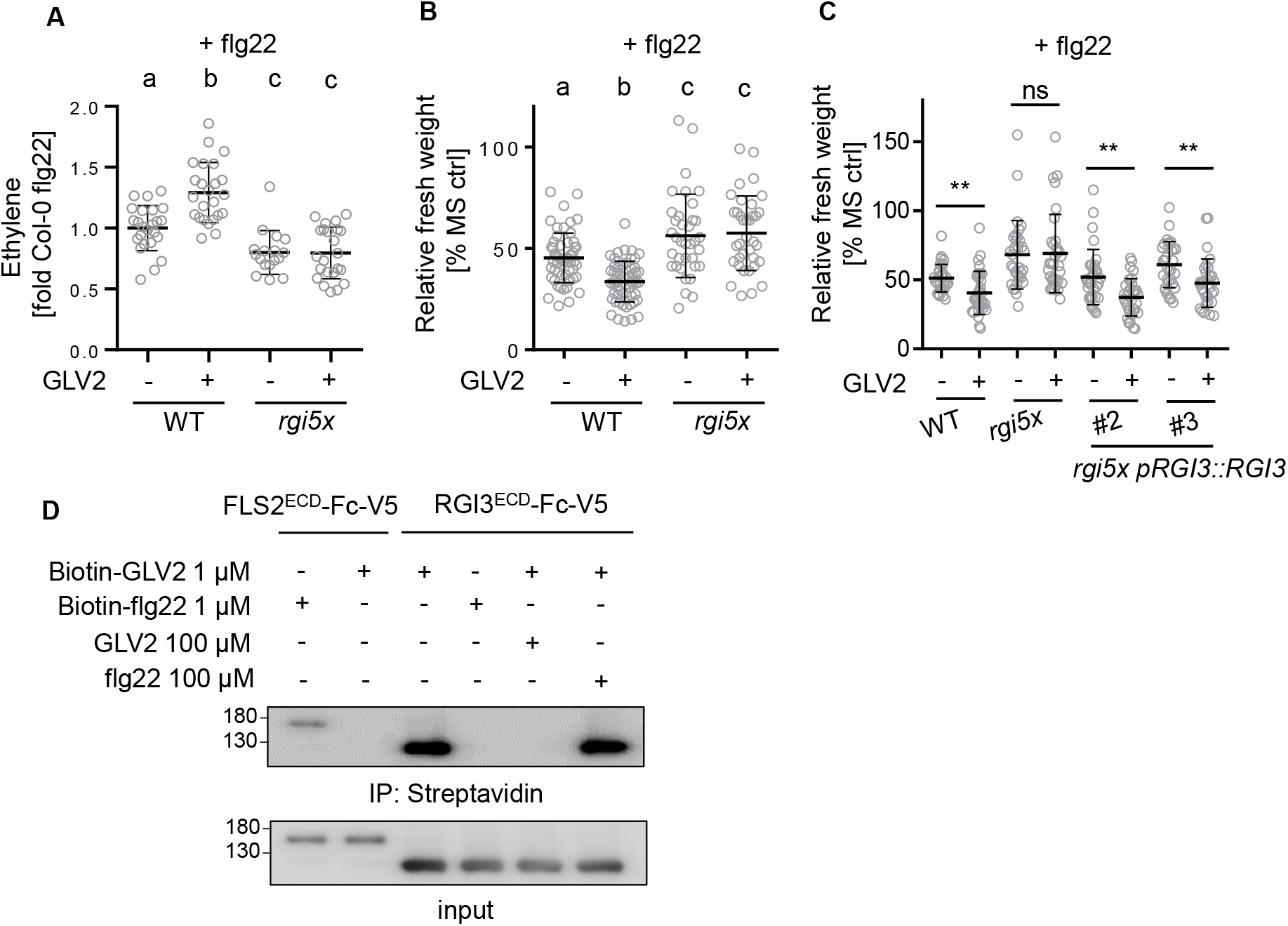
RGI3 is required for GLV2 perception during FLS2-triggered PTI. (A) Ethylene production was measured in WT and *rgi5x* 3.5 h after elicitation with 500 nM flg22 or co-treatment with 1 μM GLV2. Shown is the mean of n=17-24 from 4 independent biological replicates −/+ SD (one-way ANOVA, Tukey post-hoc test, a-b p<0.001; a-c, p<0.05). Ethylene production was normalized to the average of WT responses to flg22 set as 1. The enhancing effect of GLV2 on flg22-induced ethylene accumulation is abolished in *rgi5x*. (B) 5-day-old seedlings of WT and *rgi5x* were treated with 10 nM flg22 or co-treated with 1 μM GLV2 for 7 days before measuring fresh weight. Shown is the mean of relative fresh weight compared to the MS medium control −/+ SD, n=36-57 pooled from 4 independent biological repeats (one-way ANOVA, Tukey post-hoc test, a-b p<0.001; a-c, p<0.01). GLV2 does not increase flg22-induced SGI in *rgi5x*. (C) 5-day-old seedlings of WT, *rgi5x* and *rgi5x pRGI3*::*RGI3 #2* and *#3* were treated with 10 nM flg22 or co-treated with 1 μM GLV2 for 7 days before measuring fresh weight. Shown is the mean of relative fresh weight compared to the MS medium control −/+ SD, n=32 pooled from four independent biological repeats (Students T-test, **p<0.01). Native promoter-driven expression of RGI3 can complement the defects in GLV2 sensing of the *rgi5x* mutant (D) Streptavidin pulldown of RGI3^ECD^ and FLS2^ECD^ with 1 μM Biotin-GLV2 or 1 μM Biotin-flg22. Binding competition was performed with a 100-fold excess of unlabelled flg22 or GLV2. For protein detection, western blots were probed with α-V5 antibodies. Biotin-GLV2 directly and specifically interacts with RGI3^ECD^. Similar results were obtained in 3 independent experiments.

### RGI3 forms a flg22-induced complex with FLS2

Phytocytokines can regulate immune responses mediated by PRRs, but the mechanistic details of their mode-of-action remains largely unknown (Segonzac and Monaghan, 2019). Here, we found that GLV2 peptides can promote FLS2-triggered immunity. Thus, we set out to understand the potential molecular mechanisms employed by the GLV2-RGI3 module for the modulation of FLS2 function. Hence, we tested, whether RGI3 may be part of a FLS2 receptor complex. We expressed RGI3-GFP in *Nicotiana benthamiana* leaves and performed co-immunoprecipitation experiments using the LRR-RK CLAVATA1 (CLV1) as a negative control. FLS2-HA weakly associated with RGI3-GFP but not with CLV1-GFP (Figure 5). Interestingly, the RGI3-FLS2 interaction was strongly enhanced after flg22 treatment, suggesting that FLS2 forms an flg22-induced complex with RGI3. This indicates that RGI3 can be part of PRR complexes thereby highlighting a novel function for this family of RKs.

**Figure 5:**
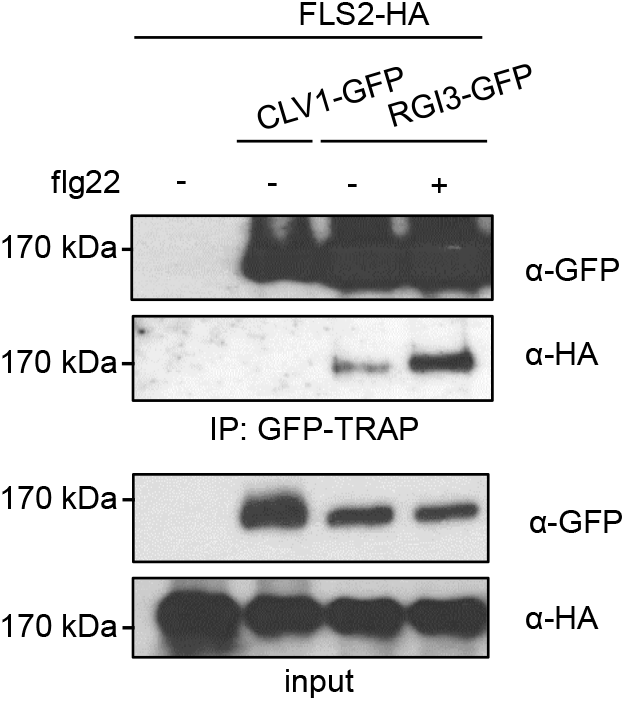
RGI3 forms a flg22-induced complex with FLS2. Co-immunoprecipitation experiment upon co-expression of FLS2-HA with either CLV1-GFP or RGI3-GFP in *N. benthamiana*. Protein extraction was performed 30 min after mock or 1 μM flg22 treatment. IP was done using GFP-TRAP agarose beads. Western blots for protein detection were probed with α-GFP or α-HA antibodies. FLS2-HA specifically interacts with RGI3-GFP and the association is enhanced after flg22 treatment. Similar results were obtained in 3 independent biological repeats.

### GLV2 perception by RGIs regulates posttranscriptional FLS2 abundance

GLV signalling was shown to control the stability of the PLT transcription factor and the auxin efflux carrier PIN2 during root stem cell niche maintenance and root gravitropism, respectively (Matsuzaki et al., 2010; Whitford et al., 2012). We tested whether RGIs may similarly control the accumulation of FLS2. Indeed, we found that both treatment of seedlings with GLV2 or *GLV2* precursor overexpression resulted in increased FLS2 protein levels (Figure 6A). In contrast, the abundance of the FLS2 co-receptor BRASSINOSTEROID INSENSITIVE 1-ASSOCIATED RECEPTOR KINASE 1 (BAK1) was not altered by GLV2 treatment or *GLV2* overexpression, suggesting that GLV2 modulates specifically FLS2 protein levels. Because *FLS2* transcript accumulation was not affected by GLV2 treatment (Figure 6B), we propose that GLV2 signalling modulates FLS2 accumulation at the posttranscriptional level. This is reminiscent of the role of GLV1 and GLV3 in increasing the abundance of PIN2 during root gravitropism (Whitford et al., 2012). Consistent with this newly discovered function for the GLV2 peptide, the *rgi5x* mutant showed impaired FLS2 accumulation (Figure 6C). Collectively, our results suggest that FLS2 abundance is controlled by GLV2-RGI signalling. These findings reveal a novel function and mechanism of GLV phytocytokines in plant immunity.

**Figure 6:**
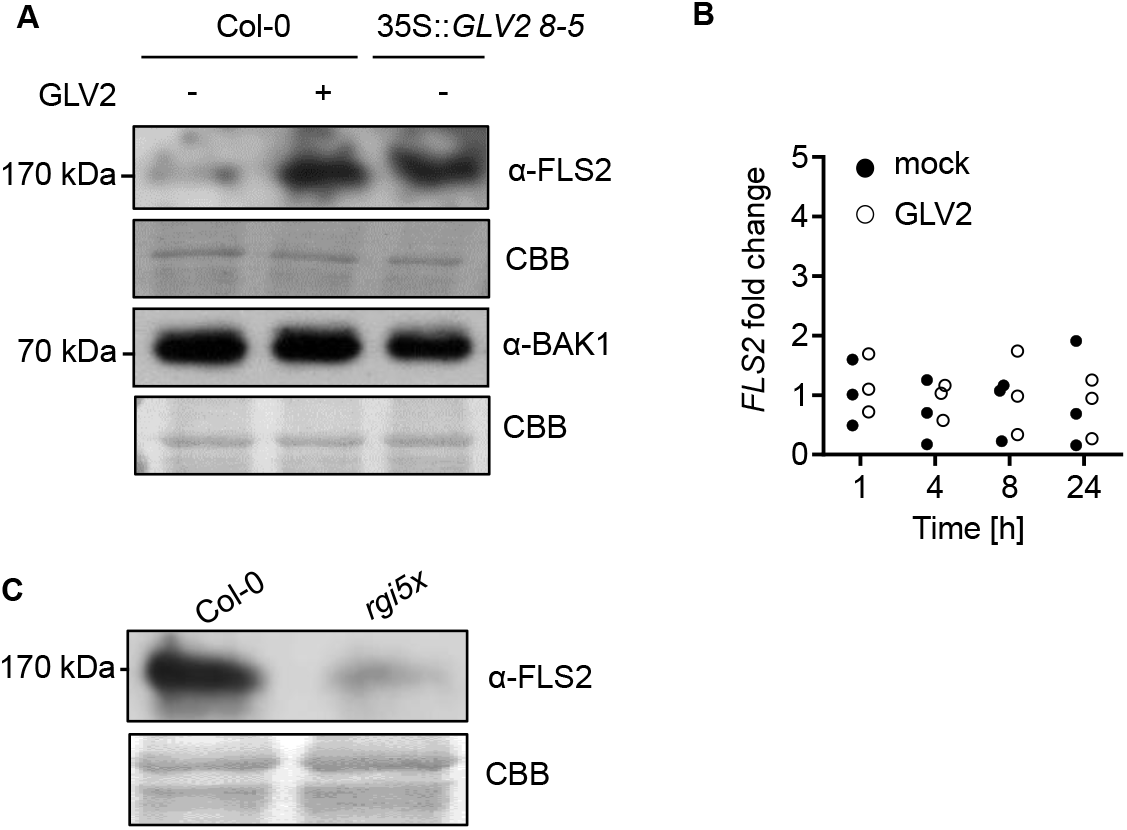
GLV2 perception by RGIs promotes maximal FLS2 abundance independent of transcription. (A) 12-day-old WT or *35S::GLV2 8-5* seedlings were treated with mock or 1 μM GLV2 for 24h before protein extraction. Western blots were probed with α-FLS2 or α-BAK1 antibodies. CBB: Coomassie brilliant blue. GLV2 treatment and precursor overexpression results in increased FLS2 protein levels. (B) Quantitative real-time PCR of *FLS2* transcripts from 12-day-old seedlings treated with mock or 1 μM GLV2 for the indicated time, n=3 biological replicates. *UBQ* was used as a house keeping gene for normalization. *FLS2* transcript levels did not change after GLV2 treatment (C) Protein extraction was performed with 12-day-old WT or *rgi5x* seedlings. Western blot was probed with α-FLS2 antibodies. The *rgi5x* mutant shows reduced FLS2 protein levels. Similar results were obtained in 3 independent experiments.

## Discussion

Tight regulation of the on and off set of immune responses is critical to ensure the plants survival and to prevent excessive defence responses in the absence or ceasing of infection. The trade-off between growth and immunity can be orchestrated by the complex regulation of RK signalling through endogenous plant peptide ligands, having a dual function as growth modulators and phytocytokines (Gust et al., 2017).

GLV peptides have regulatory functions controlling root meristem activity, gravitropism and lateral root emergence but a function for this family of peptides in shoot or leaf tissue remained unknown until recently (Wang et al., 2021). We identified the mature GLV2 peptide in apoplastic wash fluids from leaves in wild type plants (Figure 1 – figure supplement 2), demonstrating that GLV2 is also expressed, processed and secreted in shoots comparable to roots (Whitford et al., 2012). Several lines of evidence show that leaf expressed and secreted GLV peptides control immunity by regulating FLS2 levels. While the overexpression of *GLV1* and *GLV2* increases flg22-triggered responses and resistance to *Pto*, the genetic elimination of leaf expressed GLVs results in impaired flg22 signalling and increased susceptibility to *Pto* (Figure 1). Treatment with excess amounts of synthetic GLV2 peptides indicates that the further processing of the mature peptide in GLV overexpression lines is likely responsible for the modulation of the FLS2 pathway (Figure 2). We show that RGIs are positive regulators of flg22-induced responses and resistance to *Pto* and that RGI3 is a GLV2 receptor in leaf tissue to mediate its function during FLS2-triggered PTI (Figure 3, 4). GLV2 promotes FLS2 protein abundance independent of transcription (Figure 6) and its receptor RGI3 forms a flg22-induced complex with FLS2 (Figure 5).

Both PIP, PEP and SCOOP signalling through RLK7, PEPR1/2 and yet elusive receptor(s) induce PTI-like response such as ROS production or upregulation of defence-related genes (Flury et al., 2013; Gully et al., 2019; Hou et al., 2014; Tintor et al., 2013). Unlike these previously described phyotocytokines, GLV2 does not induce on its own PTI-like responses but rather acts to modulate the amplitude of FLS2 immune signalling (Figure 2). Consistent with our findings, the inducible overexpression of GLV4 was shown to induce MAPK activation and expression of PTI-related genes (Wang et al., 2021). Yet, exogenous GLV4 peptide application does not elicit PTI, indicating that GLV4, as we demonstrate for GLV2 here, is rather an immune responses modulator.

PIP1 and PEP perception leads to the upregulation of *FLS2* transcription to cooperatively contribute to PTI against bacterial infection (Flury et al., 2013; Hou et al., 2014; Tintor et al., 2013). In contrast, we discovered that GLV2 signalling through RGIs increases FLS2 abundance posttranscriptionally. Thus, our study reveals a novel regulatory mechanism for phytocytokine functions independent of *FLS2* gene expression which controls optimal FLS2 protein levels for immune activation (Figure 6).

In addition to GLVs, other growth-regulatory peptides act as phytocytokines, suggesting that endogenous peptide signalling pathways linking growth to PTI may be more widespread than anticipated. Yet, the immunomodulatory molecular mechanisms connecting growth to defence pathways remain to be discovered. The PSK and PSY1 receptors PSKR1/2 and PSY1R, respectively, negatively regulate flg22-induced immune responses (Igarashi et al., 2012; Mosher et al., 2013), but it is unknown whether these are also part of PRR signalling platforms or whether their signal transduction pathways intersect indirectly downstream of receptor activation. FLS2 signalling is controlled by endogenous RALF peptides via their receptor FER which constitutively interacts with FLS2 and functions as a ligand-regulated scaffold for FLS2-BAK1 complexes (Stegmann et al., 2017). Our data now indicates that RGI3 is recruited to an flg22-activated FLS2 complex (Figure 5) providing the first line of evidence for the direct regulation of PRR signalling by a growth-related and peptide-binding RK as part of induced PRR signalling platforms.

GLV signalling was previously shown to increase the levels of specific growth-related proteins (Matsuzaki et al., 2010; Whitford et al., 2012). GLV1 and GLV3 treatment increased PIN2 stability and subcellular dynamics by elevating both intracellular PIN2-containing vesicles and PIN2 localization to the plasma membrane (Whitford et al., 2012). Similar to PIN2, FLS2 cycles constitutively between plasma membrane and Trans-Golgi network compartments (Beck et al., 2012; Men et al., 2008; Robatzek, 2007), raising the possibility that GLV2 acts via RGI3 to regulate FLS2 endomembrane dynamics. Flg22-bound FLS2 undergoes clathrin-dependent endocytosis through distinct subcellular compartments on its way to the vacuole for degradation (Beck et al., 2012; Mbengue et al., 2016). FLS2 endocytosis is potentially regulated by ubiquitination through PLANT U-BOX type E3 ubiquitin ligases (PUBs). PUB12/13 associate with the activated FLS2 complex and ubiquitinate FLS2 for subsequent degradation (Lu et al., 2011). Similarly, RGI3 is recruited to the FLS2 complex in a flg22-dependent manner (Figure 5). This raises the question whether RGI3 modulates this regulatory circuit and may directly or indirectly control posttranslational modifications underlying FLS2 stability or endomembrane dynamics. RGI4 and RGI5 contribute to antibacterial resistance, too (Wang et al., 2021). It will be important to test whether RGI4 and RGI5 also form flg22-induced complexes with FLS2 and may contribute to the sensitization of leaves to flg22.

Another scenario for the control of FLS2 levels via the RGI3-GLV pathway may be the regulation by RGI downstream factors, such as MAPKs or RITF1 (Fernandez et al., 2020; Lu et al., 2020; Shao et al., 2020; Yamada et al., 2019). Also, GLV-controlled plasma membrane protein stabilization is not a general effect as other plasma-membrane bound receptors and proteins do not change their abundance in response to GLV treatment (Whitford et al., 2012). Similarly, our work shows that BAK1 levels do not change after GLV2 application (Figure 6A). Testing known RGI signalling components for an involvement in GLV-RGI-induced PTI regulation and unravelling context-specific RGI interaction partners will shed light on the underlying mechanisms in diverse GLV-regulated processes. It will delineate whether RGIs directly modulate stability of a range of specific proteins or whether this may be executed by downstream pathway components. Also, it will elucidate if GLV-RGI recruits specific signalling partners for immune regulation or whether it is intertwined with their function in controlling growth and development.

*GLV1*/*2* gene expression and apoplastic peptide abundance is not affected by flg22 perception (Figure 1 – figure supplement 1, 2). This suggests that GLV2 is required for steady-state FLS2 availability. Auxin induces *GLV1* and *GLV2* expression (Whitford et al., 2012) raising the possibility that GLV signalling coordinates hormone responses with cell surface immunity. This may represent a mechanism to fine-tune the accumulation of FLS2 for the detection of microbial intruders under fluctuating environmental conditions. Interestingly, *GLV4* is transcriptionally upregulated after flg22 perception (Wang et al., 2021). It remains to be tested whether increased *GLV4* expression translates into more abundant peptides in the apoplast and whether GLV4-RGI4/5 similarly controls FLS2 levels or may constitute an independent pathway for danger perception. Public transcriptome data show that *GLV1* and *GLV2* expression is downregulated after infection with *Psm* and *Pto* (Figure 1-figure supplement 1). A tempting hypothesis is that bacterial pathogens actively suppress *GLV* gene expression or peptide secretion to interfere with PRR accumulation to dampen immune responses.

Plant peptides can either work cell-autonomously or travel over short to long distances for cell-to-cell communication or to coordinate response between different plant organs (Olsson et al., 2019). In the case of GLV peptides, a local function was proposed in their role as regulators of root gravitropism (Whitford et al., 2012). To address this question for leaf expressed GLVs and their influence on PTI, future studies need to dissect in which specific cell types in leaf tissues GLV peptides are expressed and whether they show overlapping expression patterns with RGIs and FLS2. It will be intriguing to analyse whether GLV2-mediated FLS2 regulation extends to roots where *FLS2* and *GLV1/2* share similar expression domains (Beck et al., 2014; Whitford et al., 2012; Zhou et al., 2020). Treatment of roots with GLV11 resulted in the transcriptional upregulation of *PROPEP1-3* genes, suggesting that GLVs may engage in a positive regulatory module for danger perception in below ground tissues (Yamada et al., 2019).

Before GLVs are secreted and exert their physiological role, they require maturation through consecutive cleavage events by SBTs on the peptide precursors’ passage through the secretory pathway. Pre-processing of GLV2 is executed by SBT6.1/S1P (Ghorbani et al., 2016; Stührwohldt et al., 2020). Intriguingly, S1P is additionally maturating RALF23 peptides which attenuate immunity by inhibiting the PRR scaffolding function of FER (Srivastava et al., 2008; Stegmann et al., 2017). Production of mature GLV peptides also requires TPST-dependent tyrosine sulfation (Kaufmann and Sauter, 2019; Song et al., 2016; Whitford et al., 2012). Similarly, GLV2^−S^ is not capable of enhancing flg22-induced immune signalling. TPST was previously shown to negatively regulate immunity against *Pto* via PSK and PSY1 peptides (Igarashi et al., 2012; Mosher et al., 2013). Our work now reveals that tyrosine-sulfated peptides can also stimulate immune responses. This highlights the complexity and interconnectivity of diverse RK-peptide pathways in the plants` adaptation to stress responses. It is remarkable how many different signalling peptides modulate immune responses and that some have opposing functions although they partially share the same enzymes for maturation. In summary, future studies need to address the complex physiological interplay of phytocytokines, including PSK, PSY1, RALF and GLV peptides, addressing the spatio-temporal expression, maturation and/or secretion dynamics of these functionally distinct peptides during immune responses.

## Materials and Methods

### Plant material and growth conditions

Arabidopsis ecotype Columbia (Col-0) was used as a wild type control for all plant assays. The CRISPR *glv* mutant plants were compared to Col-0 expressing Cas9 (Fernandez et al., 2020). The 35S::*GLV1 16-1/18-5* and 35S::*GLV2 8-5/10-4* were kindly provided by Ana Fernandez (Whitford et al., 2012). The *rgi1-1/rgi2-1/rgi3-1/rgi4-1/rgi5/1* (*rgi5x*) mutant was kindly provided by Jia Li (Ou et al., 2016). Plants for ROS burst assays, ethylene measurements and *Pto* infection experiments were grown in individual pots at 20-21 °C with an 8 h-photoperiod in environmentally controlled growth rooms. For seedling-based assays, seeds were sterilized using chlorine gas for 4 h and grown on ½ Murashige and Skoog (MS) media supplemented with vitamins, 1% sucrose and 0.8% agar at 22 °C and a 16 h photoperiod. For transient co-immunoprecipitation experiments, *Nicotiana benthamiana* plants were grown for 4 weeks at 22 °C and a 16 h photoperiod.

### Molecular cloning

To generate *pRGI3* driven *RGI3* complementation lines in the *rgi5x* background, *pRGI3::RGI3* was cloned from genomic DNA containing 1404 bp upstream of the start codon with primers pRGI3_F and pRGI3_R containing attB attachment sites for subsequent BP reactions. The PCR product was cloned into pDONR223 (Thermo Fisher, Germering, Germany) using BP clonase (Thermo Fisher). The resulting pDONR223 pRGI3::RGI3 construct was subcloned into pGWB4 (Nakagawa et al., 2007) using LR clonase (Thermo Fisher). The resulting construct was sequenced and transformed into *Agrobacterium tumefaciens* strain GV3101. Subsequently, *rgi5x* mutants were transformed via floral dip. To generate the *35S*::*RGI3*-*GFP* construct for transient expression experiments, *RGI3* was PCR amplified from Col-0 cDNA using primers RGI3_F and RGI3_R. The PCR product was cloned into pDONR223 (Thermo Fisher) and subsequently recombined with pK7FWG2 (VIB Ghent) using LR clonase (Thermo Fisher).

To generate fusion constructs for transient overexpression of CLV1-GFP and FLS2-HA, full-length coding sequence of *CLV1* was PCR amplified from Col-0 cDNA using primers CLV1_F and CLV1_R and cloned into a GoldenGate-adapted pUC18-based vector similar to previously described (Engler et al., 2008). CLV1 was further domesticated to remove an internal AarI restriction site via whole-plasmid PCR amplification using primers CLV1-AarI_F and CLV1-AarI_R and GoldenGate-based plasmid recirculation. Full length coding sequence of *FLS2* was cloned similarly. Two fragments were amplified to remove an internal EciI restriction site with primers FLS2_1_F, FLS2_1_R (FLS2 fragment 1) and FLS2_2_F, FLS2_2_R (FLS2 fragment 2). An internal *FLS2* AarI restriction site was removed as described above using primers FLS2-AarI_F and FLS2-AarI_R. *CLV1* and *FLS2* coding sequences were subsequently fused to C-terminal meGFP or 1xHA epitope tags with a ten- or five-glycine linker, respectively, and CaMV35S promoter and terminator sequences. Finally, full expression cassettes were cloned into a GoldenGate-adapted binary pCB302 vector. Sequences of all primers can be found in Supplementary Table 1.

### ROS burst measurements

For ROS burst measurements, leaf discs (3 mm diameter) of Arabidopsis were collected in 96 well plates using biopsy punchers and floated overnight on 100 μl H_2_O in a 96-well plate. Elicitation was performed with the indicated concentration of peptides and 2 μg/ml horseradish peroxidase (Type II, Roche, Penzberg, Germany) and 5μM L-012 (FUJIFILM Wako chemicals, Neuss, Germany). Luminescence was measured as relative light units (RLU) in 1min intervals using a Tecan F200 luminometer (Tecan Group Ltd., Männedorf, Switzerland).

### Pathogen inoculation experiments

*Pto* DC3000 COR-bacteria were streaked out from glycerol stock on fresh King’s B media plates containing 1% agar and 50 μg/mL rifampicin and 50 μg/mL kanamycin. Bacteria were collected from plates with a sterile pipette tip and resuspended in water containing 0.04% Silwet L77 (Sigma Aldrich, St. Louis, USA) to an OD_600_ = 0.2 (10^8^ cfu/mL). The bacterial suspension was sprayed on 4 to 5-week-old plants, which were subsequently covered with lids for 3 days. 3 leaf discs per sample from different plants were collected in microfuge tubes and ground with a tissue lyser (Qiagen, Düsseldorf, Germany). Serial dilutions were plated on LB agar before counting colonies. For inducing resistance with GLV2 peptides, 4-to-5-week-old plants were infiltrated with 1 μM GLV2 and incubated for 24 h. Subsequently, a suspension of *Pseudomonas* syringae pv. *tomato* DC3000 bacteria was prepared as above to an OD_600_ = 0.0002 (10^5^ colony forming units per mL) and syringe-infiltrated into pre-treated leaves. Two days after inoculation, samples were collected as described above.

### Ethylene measurements

Arabidopsis leaf discs (3 mm diameter) were floated overnight in H_2_O containing petri dishes. Three leaf discs were transferred to a glass vial (6ml volume) with 500 μl H_2_O before adding flg22 and/or GLV2 to a final concentration of 500 nM and 1 μM, respectively. Glass vials were tightly sealed with a rubber lid and incubated under slight agitation for 3.5 h on a horizontal shaker. 1 ml air was extracted with a syringe and injected into a gas chromatograph to measure ethylene.

### RNA isolation and quantitative RT-PCR

Four 2-week-old seedlings grown in liquid MS were treated with 100 nM flg22 and/or 1 μM GLV2 for 24 h and ground in liquid N_2_. Total RNA was extracted using Direct-zol™ RNA Miniprep Plus kit (Zymo Research, Freiburg, Germany) according to the manufacturer`s instructions. cDNA synthesis was performed with 2 μg RNA per sample using RevertAid First Strand cDNA Synthesis kit (Thermo Fisher). *PR1* expression was detected using primers qPR1_F and qPR1_R by quantitative real-time PCR using Maxima SYBR green mix (Thermo Fisher) and the AriaMx Real-Time PCR system (Agilent Technologies, Santa Clara, USA). The house keeping gene *UBQ* was used as a reference gene using primers qUBQ_F and qUBQ_R. *FLS2* transcript levels were determined using primers qFLS2_F and qFLS2_R. To detect *RGI1*-*5* transcript levels in mature leaves of WT and *rgi5x* and *RGI3* expression in *pRGI3::RGI3* lines primers semi-qRT-RGI1-5_F and R were used. Sequences of all primers can be found in Supplementary Table 1.

### Root growth and seedling growth inhibition

Seeds were surface-sterilized and grown on MS Agar plates for 5 days before transferring individual seedlings in each well of a 48-well plate containing MS medium with 1 μM GLV2 and/or flg22 in the indicated concentrations before measuring fresh weight 7 days after transfer. To determine root length in *rgi5x* and *rgi5x pRGI3*::*RGI3* lines, seeds were grown vertically on MS Agar plates for 7 days before measuring root length

### Binding of GLV2 to RGI3

The ECD constructs of FLS2 and RGI3 were previously described (Smakowska-Luzan et al., 2018) and expressed via transient transfection of *Drosophila* S2 cells at 27 °C. Upon transfection using Effectene (Qiagen), the temperature was changed to 21 °C. 1 mM CuSO_4_ for induction of protein was added to the culture 24 h after transfection. The supernatant was collected 3 days after induction. Protease inhibitors (Roche) and 0.02 % Sodium azide were added to the supernatant containing the recombinant ECD and then stored at 4 °C before use. The supernatants were incubated with biotin-labeled peptides over-night. For competition assays, unlabelled peptide was added 20 min prior to biotin-labelled peptide. The next day, the supernatant was incubated with PBS equilibrated Streptavidin conjugated agarose resin (Thermo scientific) for 2 h at 4 °C. After 3 times washing of the resin with PBS containing 1% NP-40, bound proteins were eluted with SDS sample buffer, and then eluted proteins were subjected to immunoblotting using anti-V5 antibodies (Thermo Scientific).

### Protein extraction and co-immunoprecipitation experiments

To determine FLS2 and BAK1 levels, seedlings were grown for 5 days before transferring 2-3 seedlings in each well of a 24-well plate containing MS medium for 7 days. Afterwards, MS medium was replaced with fresh MS medium with or without 1 μM GLV2. 24 h later, seedlings were harvested in liquid N_2_ and subjected to protein extraction (50 mM Tris-HCl pH 7.5, 50 mM NaCl, 10% glycerol, 2 mM Ethylenediaminetetraacetic acid, 5 mM dithiothreitol, 1% protease inhibitor cocktail (Sigma Aldrich), 1 mM phenylmethanesulfonyl fluoride and 1 % IGEPAL). After SDS PAGE and western blot, proteins were detected using α-FLS2 and α-BAK1 antibodies (Chinchilla et al., 2007).

For co-immunoprecipitation experiments upon transient expression in *N. benthamiana*, 4-5 week-old leaves were infiltrated with Agrobacteria containing constructs for 35S::FLS2-HA, 35S::CLV1-GFP and/or 35S::RGI3-GFP expression. In all cases constructs were co-infiltrated with a P19 silencing suppressor construct. Leaves were harvested 2 days post-infiltration and treated with 1 μM flg22 for 30 min via vacuum infiltration before freezing in liquid N_2_. Protein extraction was performed with 50 mM Tris-HCl pH 7.5, 100 mM NaCl, 10% glycerol, 2 mM Ethylenediaminetetraacetic acid, 5 mM dithiothreitol, 1% protease inhibitor cocktail (Sigma Aldrich), 1 mM phenylmethanesulfonyl fluoride and 0.5% IGEPAL. Immunoprecipitation was performed with magnetic GFP-TRAP beads (Chromotek, Martinsried, Germany). After SDS PAGE and western blot, proteins were detected using α-HA (Chromotek) and α-GFP antibodies (Santa Cruz Biotechnology, Dallas, USA).

### Detection of GLV2 peptides from apoplastic wash fluids using targeted proteomics

Preparations of apoplastic wash fluids were performed as previously described (Nakano et al., 2020). In brief, 4-5-week-old soil-grown plants were detached from root tissue and submerged in apoplastic wash fluid buffer (5 mM NaC_2_H_3_O_2_^−^, 0.2 M CaCl_2_, pH 4.3) and vacuum infiltrated. Upon centrifugation, eluates were collected and subjected to chlorophenol extraction according to previous protocols (Ohyama et al., 2008). After precipitation, peptides were dissolved in H_2_O and analysed by targeted mass spectrometry using Parallel Reaction Monitoring (PRM). Synthetic GLV2 reference peptide (Supplementary Table 2) was used for PRM assay setup and optimization. PRM measurements were performed with a 50-min linear gradient on a Dionex Ultimate 3000 RSLCnano system coupled to a Q-Exactive HF-X mass spectrometer (Thermo Fisher Scientific). The mass spectrometer was operated in PRM and positive ionization mode. MS1 spectra (360–1300 m/z) were recorded at a resolution of 60,000 using an AGC target value of 3×10^6^ and a MaxIT of 100 ms. Targeted MS2 spectra were acquired at 60,000 resolution with a fixed first mass of 100 m/z, after higher-energy collisional dissociation (HCD) with 26% NCE, an AGC target value of 1×10^6^, a MaxIT of 118 ms and an isolation window of 1.3 m/z. Within a single PRM run 17 peptide precursors were targeted (the GLV2 target peptide in several charge states as well as 14 retention time reference peptides) without retention time scheduling. The cycle time was ~2.1 s, which lead to ~10 data points per chromatographic peak. PRM data analysis was carried out using the software tool Skyline (version 64-bit 20.2.0.286, (Maclean et al., 2010)). Interferences and peak integration boundaries were reviewed manually, considering the ten most intense transitions (precursor-fragment ion pairs) of the GLV2 peptide. The total GLV2 peptide intensity was computed by summing up all transition intensities.

### Synthetic peptides and elicitors

The flg22 peptide was kindly provided by Dr. Justin Lee (IPB Halle). GLV2 and GLV^−S^ peptides were synthesized by Pepmic (Suzhou, China) to a purity of minimum 90%. The Biotin-flg22 peptide was synthesized in house at the Gregor Mendel Institute. All peptides were dissolved in H_2_O. Sequences of all peptides used in this study can be found in Supplementary Table 2.

### Statistical analysis

Analysis was carried out using the GraphPad Prism software (San Diego, USA). The detailed statistical method employed is provided in the respective figure legends.

## Data availability

All targeted proteomic raw data and the Skyline analysis file have been deposited to Panorama Public (Sharma et al., 2018) and can be accessed under https://panoramaweb.org/GLV2_peptide.url (ProteomeXchange ID: PXD023855).

## Acknowledgements

The work was supported by core funds from the Technical University of Munich (Martin Stegmann, Patricia Zecua-Ramirez, Ralph Hückelhoven, Christina Ludwig). Ho-Seok Lee and Youssef Belkhadir are supported by grants from the Austrian Academy of Science through the Gregor Mendel Institute and by the Austrian Science Fund (FWF) (I 3654). Zachary Nimchuk and Brenda Peterson are supported by a National Science Foundation Plant Genome Research Program Grant (Z.L.N., no. PGRP-1841917). We thank Lars Raasch (Chair of Phytopathology, Technical University of Munich) for cloning and providing the constructs for transient overexpression of FLS2-HA and CLV1-GFP in *N. benthamiana*. We also thank Franziska Hackbarth, Hermine Kienberger, Lara Wanner and Verena Breitner for technical assistance at BayBioMS, as well as Miriam Abele for mass spectrometric support.

## Author contribution

Martin Stegmann planned the project. Martin Stegmann, Youssef Belkhadir, Christina Ludwig and Ralph Hückelhoven designed and conceived the experiments and analysed data. Zachary L. Nimchuk oversaw work. Martin Stegmann, Patricia Zecua-Ramirez, Christina Ludwig, Ho-Seok Lee and Brenda Peterson performed the experiments. Martin Stegmann wrote the manuscript with input from all the authors.

## Supplementary figures

**Figure 1 – figure supplement 1:**
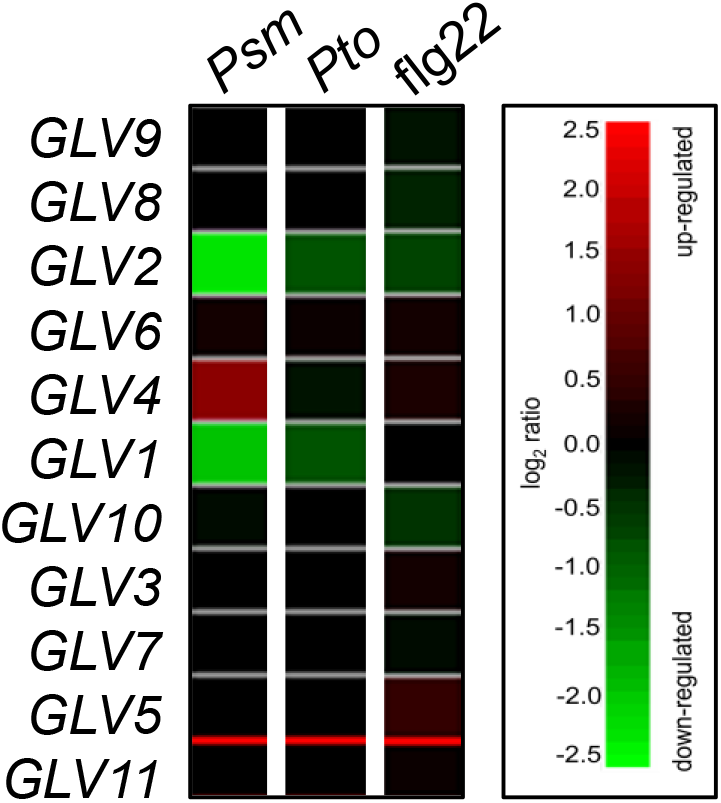
*GLV* peptides are differentially expressed in response to *Psm* infection and involved in flg22-triggered PTI. Data was obtained using Genevestigator software and are based on the AT_nRNASeq_ARABI_GL-1 data set. Both *GLV1* and *GLV2* are transcriptionally downregulated after infection with *Psm* and to a lesser extent after infection with *Pto*. Treatment with flg22 does not affect *GLV* gene expression.

**Figure 1 – figure supplement 2:**
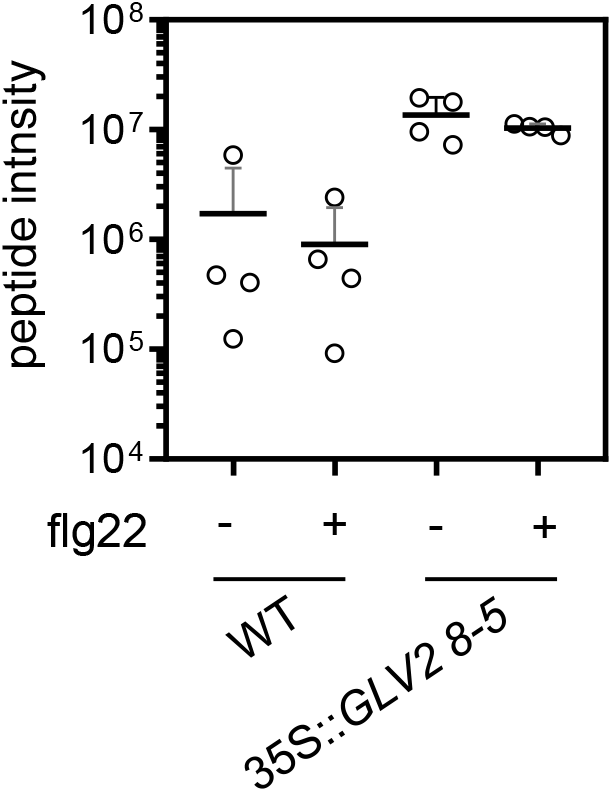
GLV2 peptide is detectable in apoplastic wash fluids. Targeted proteomics (Parallel Reaction Monitoring, PRM) detection of the GLV2 peptide (DMDY[+80]NSANKKRP[+16]IHN) from apoplastic wash fluids after treatment with 1h mock or 100 nM flg22. An optimized targeted proteomics assay was setup using a synthetic reference GLV2 peptide (Supplementary Figureary Table 2). Shown are label-free mean peptide intensities. The standard deviation is indicated from intensities +SD, n=4 biological replicates.

**Figure 1 – figure supplement 3:**
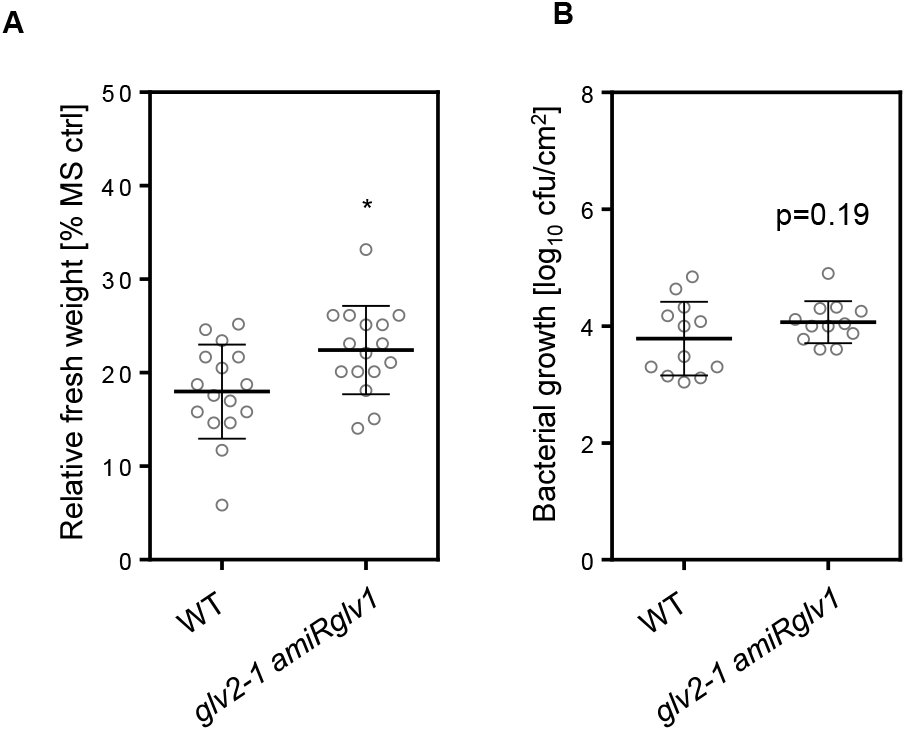
Loss of *GLV1* and *GLV2* shows moderate reduction in flg22-triggered PTI. (A) 5 day-old seedlings of the indicated genotypes were treated with 100 nM flg22 for 7 days before measuring fresh weight. Shown is the mean of relative fresh weight compared to the MS control −/+ SD, n=16 (Students T-test, *p<0.05). The *glv2-1 amiRglv1* lines show reduced flg22-triggered seedling growth inhibition. (B) Colony-forming units (cfu) of *Pto* DC3000 COR-four days after spray infection of the indicated genotypes; n=12 from 3 pooled experiments −/+ SD. Students T-test indicates p=0.19. The *glv2-1 amiRglv1* lines do not show a significantly altered infection phenotype. Only a trend towards enhanced suscpetibility was observed.

**Figure 2 – figure supplement 1:**
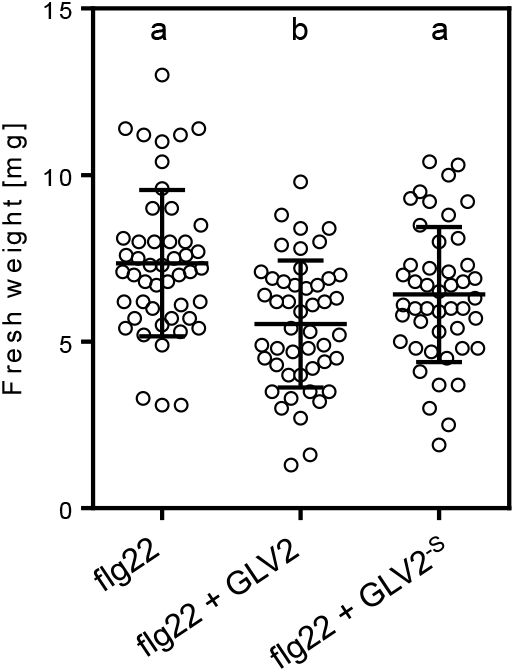
GLV2 tyrosine sulfation is required for its modulatory function on flg22 responses. (A) 5-day-old WT seedlings were treated with 10 nM flg22 or co-treated with 1 μM GLV2 or 1 μM GLV2^−S^ for 7 days before measuring fresh weight. Shown is the mean of relative fresh weight compared to the MS medium control −/+ SD, n=45-47 from four pooled biological replicates (one-way ANOVA, a-b p<0.001). The GLV2^−S^ mutant peptide does not significantly enhance flg22-induced seedling growth inhibition.

**Figure 3 – figure supplement 1:**
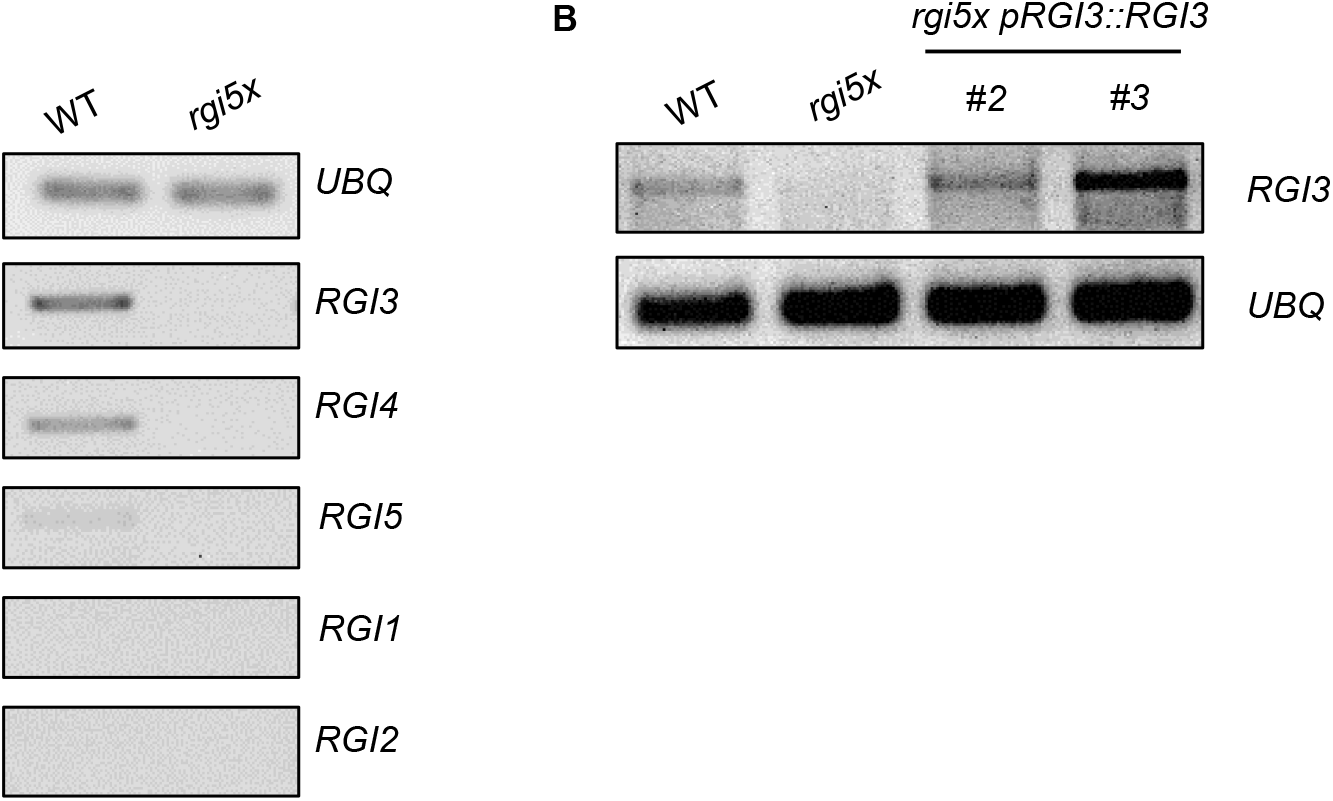
Characterization of *RGI* expression in mature leaves and *RGI3* expression in *rgi5x pRGI3*::*RGI3*. (A) Semiquantitative reverse transcription PCR showing expression of *RGI1-5* in 5–6-week-old leaves of WT and *rgi5x*. *UBQ* was used as a control. *RGI3* and *RGI4* show comparable expression levels. *RGI5* shows weaker transcript levels and *RGI1* and *RGI2* expression was not detectable in mature leaves. (B) Semiquantitative reverse transcription PCR showing expression of *RGI3* in transgenic *rgi5x pRGI3::RGI3* lines. *UBQ* was used as a control. Both lines of *rgi5x pRGI3*::*RGI3* show *RGI3* expression in mature leaves.

**Figure 3 – figure supplement 2:**
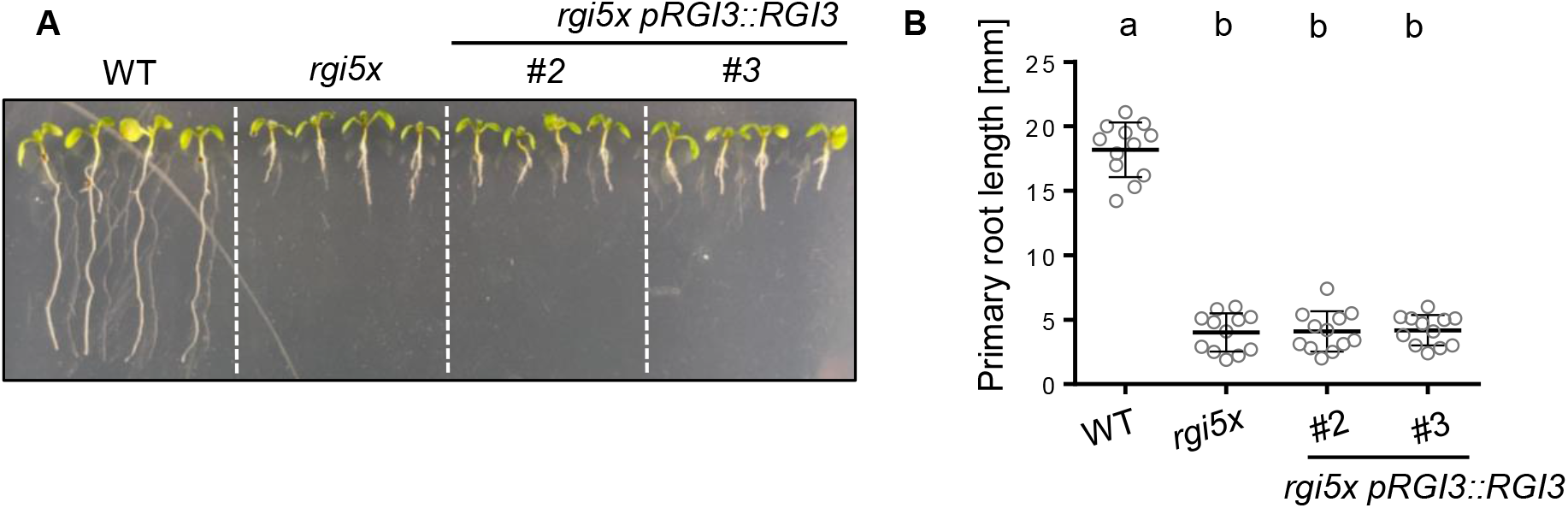
*RGI3* does not complement the root growth defect of *rgi5x*. (A) Representative pictures taken 7 days after vertical germination of the indicated genotypes on MS Agar plates. (B) Quantification of main root length of seedlings shown in A. Shown is the mean of n=12 +/− SD (one-way ANOVA, Tukey post-hoc test, a-b p<0.001). Expression of *pRGI3::RGI3* does not complement the short root phenotype of the *rgi5x* mutant.

## Supplementary Tables

**Supplementary Table 1:**
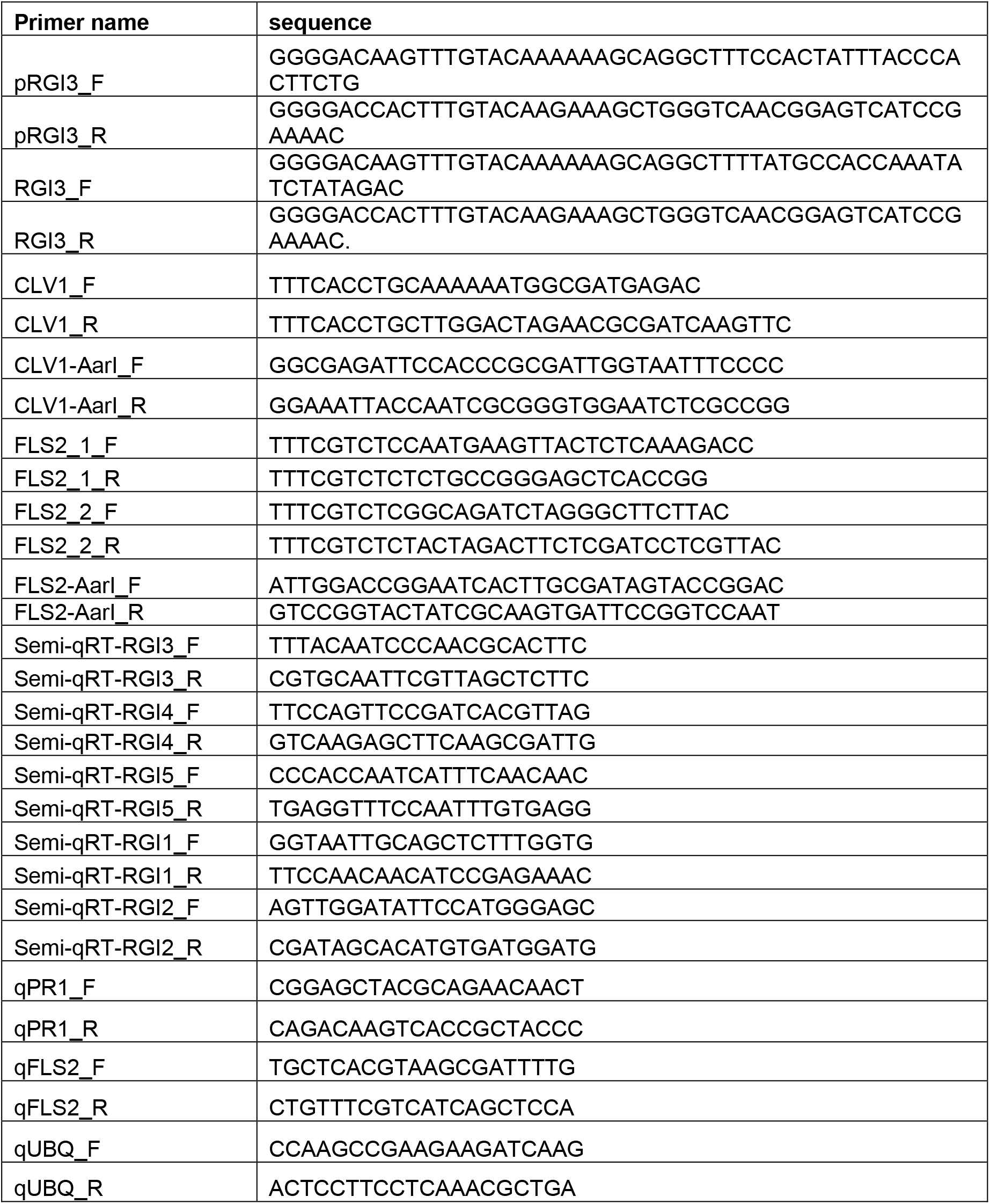
Primers used in this study.

**Supplementary Table 2:**
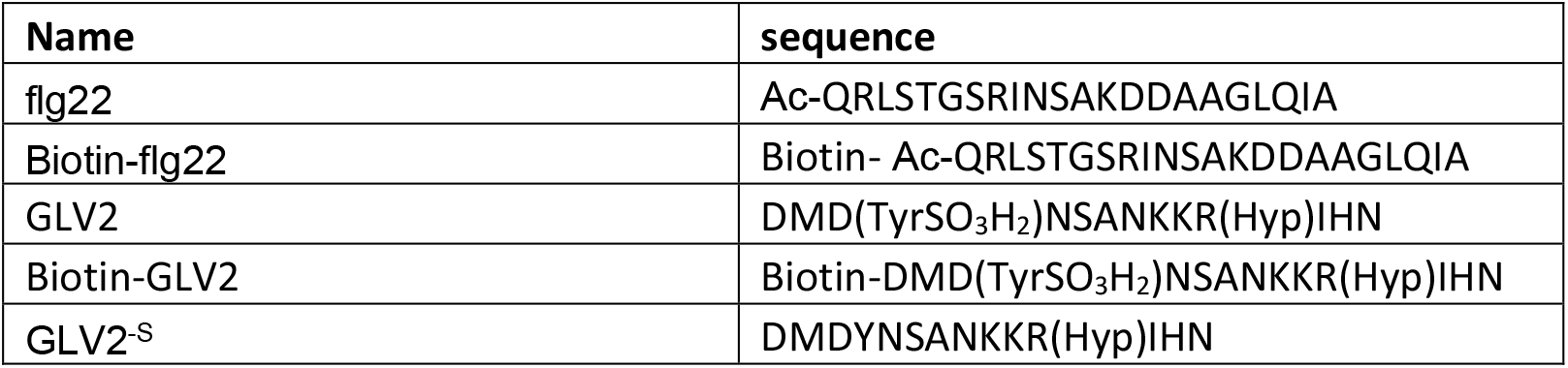
Sequences of peptides used in this study.

